# Symbiont-specific uptake is mediated by integrins in cnidarian larvae

**DOI:** 10.1101/2025.01.21.633834

**Authors:** Victor A. S. Jones, Melanie Dörr, Isabelle Siemers, Sebastian Rupp, Joachim M. Surm, Ira Maegele, Sebastian G. Gornik, Meghan Ferguson, Annika Guse

## Abstract

The symbiotic relationship between dinoflagellate algae and their cnidarian hosts is fundamental to the health of coral reefs. The selection of appropriate symbionts is paramount for the host to gain valuable nutrients and could be tailored to increase stress tolerance against anthropogenic induced changes in ocean environments, such as coral bleaching in response to ocean warming. Previous research suggests glycan-lectin interactions play a role in symbiont uptake; however, blockage of such interactions does not fully inhibit symbiosis establishment, suggesting other receptors are at play. Potential candidates include RGD peptide binding integrins, which are known to mediate phagocytosis of microbes in other systems. Here, we used a combination of cnidarian model systems and human cell lines to determine if integrins facilitate symbiont recognition and uptake. Integrins are highly expressed in the endodermal tissue of the host, where symbiosis takes place, and upon uptake into endodermal cells, symbionts altered the expression of integrins and downstream signaling molecules. Blockage of integrin binding sites with RGD competitor peptides reduced symbiont uptake, but had no effect on the general uptake of non-symbiotic algae, or uptake in a non-symbiotic cnidarian. In addition, inert beads coated with integrin RGD peptide ligands were phagocytosed more readily than beads coated with scrambled peptide. Finally, overexpression of RGD-binding integrins in human cells increased symbiont uptake and mutation of the active binding site abolished uptake. Our findings reveal RGD-binding integrins as key players in symbiosis establishment and shed light on the evolutionary functions of integrins as phagocytic receptors.

**Significance statement:** Corals engage in a symbiotic partnership with photosynthetic algae to survive in challenging environments. To date it is largely unclear how the two partners recognize each other. Using a comparative model systems approach, we have identified evolutionary conserved integrins as molecular receptors for specifically engulfing symbionts, but not other algae. This suggests that integrins allow the host to distinguish between symbiotic and non-symbiotic algae and preferentially take up symbionts. Our findings establish a new paradigm for symbiosis establishment in corals and shed light on the ancient function of integrins as environmental sensors.

## Introduction

To increase the chances of survival in competitive oligotrophic environments, some organisms have evolved ways to work in cooperation to mutually thrive. In shallow nutrient-poor waters across the tropics, reef-building corals have acquired the ability to establish a mutualistic symbiotic lifestyle with dinoflagellate algae of the family Symbiodiniaceae (1, 2). Using bidirectional nutrient transfer, these endosymbiotic dinoflagellates, located inside host endodermal cells, provide over 90% of the host’s nutritional needs in the form of photosynthetically fixed carbon, while the algae receive inorganic nutrients and shelter from herbivores (3–6). The success of coral reef ecosystems is strictly reliant on this fundamental association and is greatly threatened by the anthropogenic climate crisis, causing increased rates of symbiosis breakdown and subsequent coral bleaching (7–9).

If not challenged by extensive stress, host-dinoflagellate associations are stable over the lifetime of an individual host, however, most coral species produce aposymbiotic (symbiont-free) larvae that take up algae from their surroundings through horizontal (environmental) transmission (10). In this way, endosymbiosis must be re-established each generation, allowing corals to acquire symbionts specifically adapted to the local environments in which they settle (11–13). However, considering the broad and diverse abundance of microorganisms present during symbiosis onset, decisive mechanisms must exist to ensure symbiosis is established with the desired partner.

A series of complex steps, sometimes termed “winnowing,” have been proposed to facilitate symbiosis onset in various symbiotic systems, all of which are required to establish a stable association with a suitable partner (14). These steps span symbiont recognition, phagocytosis, long-term immune evasion, and maintenance (15). Genomic and cellular studies provide growing evidence that the first steps of winnowing likely resemble microbial invasion of animal and plant hosts (14, 15). Thus, in light of host-microbe interactions governed by pattern recognition receptors (PRRs) on the host plasma membrane and microbe-associated molecular patterns (MAMPs) on the surface of microbial targets, it is a widespread idea that symbiont phagocytosis is receptor-mediated and specific (16). However, it has yet to be determined which receptors or signaling pathways would be involved in symbiont-specific receptor-mediated phagocytosis since multiple molecular mechanisms have been proposed.

Lectin-glycan interactions are among the best-studied pairings involved in inter-partner recognition. They are common surface molecule MAMP-PRR interactions in innate immune responses and mutualistic endosymbiosis, and they have been extensively explored in the context of coral-algal symbiosis (17–19). Various glycan-binding lectins are present in corals (20–23) and have been shown to detect glycans on the symbiont surface (24–26). Surface glycan profiles differ among multiple symbiont clades and strains, potentially providing a basis for symbiont-specific selection (25, 27, 28). Research on the role of lectin-glycan interactions in coral-algal symbiosis are mostly based on enzymatic cleavage of glycan residues from symbionts, lectin addition to mask symbiont glycans, or lectin antibodies and glycans to competitively block host lectin-binding sites (21, 23, 26, 29, 30). However, broad digestion of cell surface proteins using trypsin led to a significant decrease in symbiont uptake in *Fungia scutaria* larvae, while digestion with glycan-specific N-glycosidase had little effect on symbiont uptake (26). Additionally, infection is not always impaired when symbiont surface glycans are masked with exogenous lectins, and surface glycan profiles were shown to differ only subtly between compatible and incompatible symbiont strains, providing no foundation to explain species-specific host colonization rates (31). These observations indicate that additional mechanisms for symbiont-specific identification and uptake exist in cnidarians.

Considering that other factors and receptors have been described to contribute to symbiont phagocytosis, such as complement and scavenger receptors (26, 29, 32–34), it appears that no singular recognition process governs the first step of winnowing and an interplay between various recognition pathways seems plausible.

Promising candidates for PRR’s involved in symbiont recognition are integrins. Integrins are heterodimeric transmembrane cell-surface receptors that contain non-covalently associated alpha and beta subunits. Integrins signal across the plasma membrane in both directions, “inside-out” and “outside-in”, through binding of small peptide ligands and most prominently function as receptors for cell-extracellular matrix (ECM) and cell-cell adhesion (35). Integrins and their extracellular ligands in mammals are clustered into four main classes: Arginine-Glycine-Aspartic Acid (RGD)-binding, laminin-binding, Leucine-Aspartic Acid-Valine (LDV)-binding, and collagen-binding integrins (36). Some integrins, such as RGD-binding and laminin-binding integrins, are ancient and present throughout Metazoans (35). Moreover, RGD-binding integrins are known to mediate integrin-dependent phagocytosis in other systems, such as flies and humans, making them interesting candidates for mediating symbiont phagocytosis in coral-algal symbiosis (37, 38).

In this study, we make use of a combination of experimental systems including *Exaiptasia diaphana* (commonly Aiptasia), *Acropora digitifera*, *Nematostella vectensis*, as well as human cells to address the fundamental question of how symbionts are recognized prior to uptake into cnidarian host cells (39–41). We identify Aiptasia integrins as novel receptor candidates and demonstrate that symbiont uptake, but not other algae, is enhanced by RGD-binding-integrin-mediated phagocytosis, where the binding of a symbiont-surface RGD-motif adds specificity to the efficient uptake of beneficial partners.

## Results

### Symbionts alter the expression of integrins in endodermal cells

Several pathogens rely on receptor-ligand interactions to gain entry into non-phagocytic cells, including several types of integrin interactions (42). To determine if integrins may be involved in symbiosis establishment in Aiptasia, we analyzed RNA-sequencing data from dissected endodermal cells that contained symbionts (*Breviolum sp.*, SSB01+) and compared their gene expression to endodermal cells from larvae that were exposed to, but did not contain symbionts (SSB01-) or from larvae that were never exposed to symbionts (Apo) (dataset previously published in ref 43). Endodermal cells that phagocytosed symbionts had significantly decreased expression of multiple integrin-related genes, including one integrin alpha and one beta subunit, compared to either aposymbiotic endodermal control (SSB01+ vs SSB01- and SSB01+ vs Apo, Fig. 1). Furthermore, RNA-sequencing analysis of endodermal cells containing the non-symbiotic microalgae *Microchloropsis gaditana* (Mg+), which are lost relatively rapidly after uptake yet remain intracellular long enough for RNA-sequencing (43), revealed that infection with *M. gaditana* did not induce changes in integrin gene expression (Fig. 1). These data suggest that symbionts specifically alter the expression of integrins upon uptake into endodermal cells while non-symbiotic algae do not.

**Figure 1.**
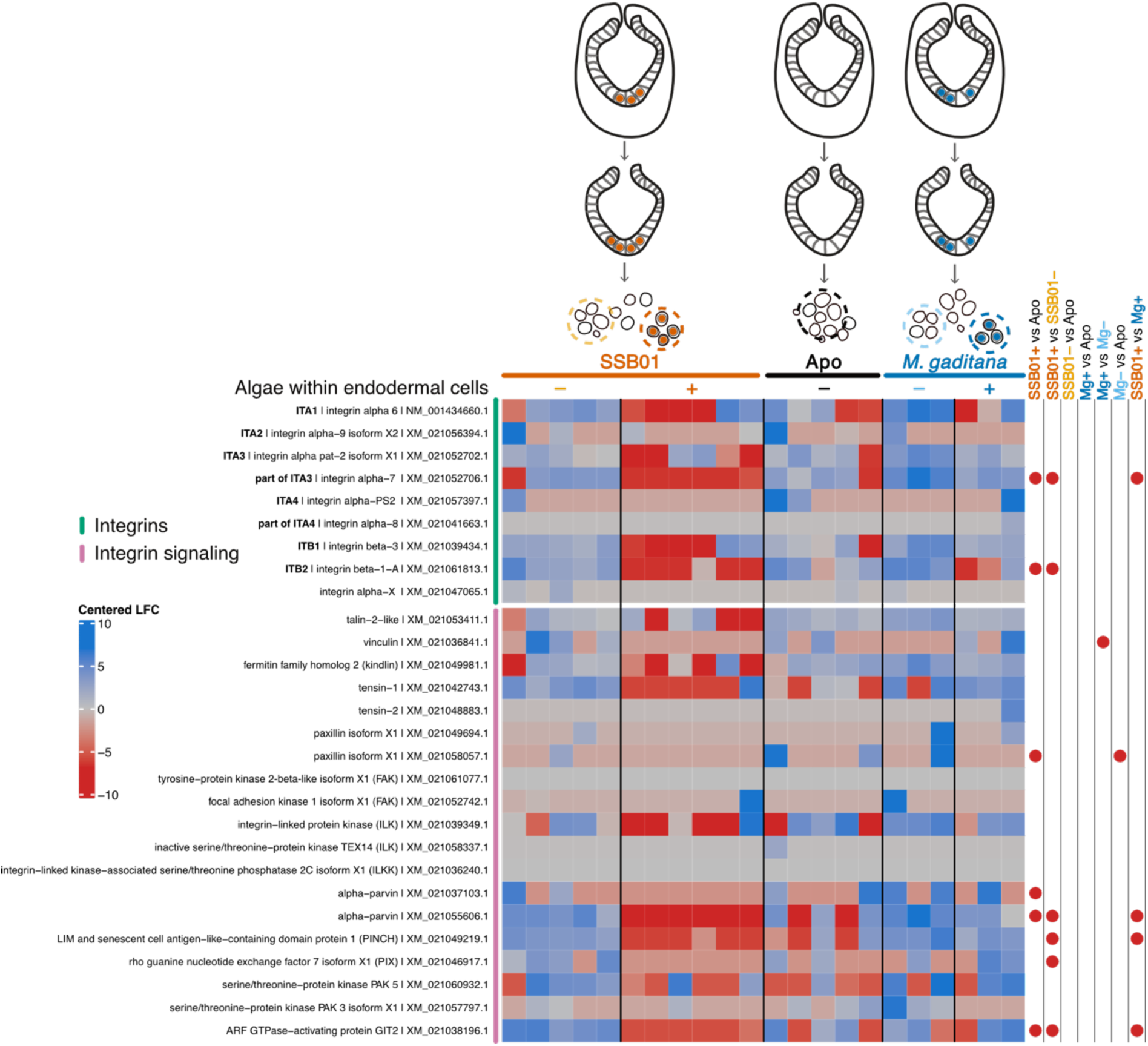
Symbionts alter the expression of integrins in endodermal cells. Aiptasia larvae exposed to either SSB01, *M. gaditana,* or left aposymbiotic for 24 hours were manually dissected to extract endodermal cells with (+) or without (-) intracellular algae. The resulting samples were processed for RNA-sequencing (43). Expression changes in integrins and their downstream signaling genes are shown. The color in the heatmap indicates the centered log_2_ fold change (LFC) according to DESeq2 (red = downregulation and blue = upregulation). Significantly differentially expressed genes (p <0.05) compared between populations of cells are indicated with red (downregulated) dots.

### Predicted RGD-binding Aiptasia integrin alpha 1 is highly expressed in larval endodermal tissue

To gain insight into the evolutionary functions of cnidarian integrins, we compared the phylogeny of alpha integrins from Aiptasia and other cnidarians to that of vertebrate integrins with known functions. Phylogenomic analysis revealed four integrin alpha subunits (ITAs)) in Aiptasia (Fig. 2A), with cnidarian ITAs falling into two cnidarian-specific clades. We propose the names cnidarian ITA Clade 1 and 2 for these. Cnidarian ITA Clade 1 contains ITA1 (NM_001434660.1, integrin alpha-6), as well as a single gene from each cnidarian species included in the analysis. Cnidarian ITA Clade 1 falls together with two other previously well-supported clades with representatives from vertebrates: PS1, which canonically bind to laminins (45); and PS2, which canonically bind to the tripeptide RGD and the related Lysin-Glycine-Aspartic Acid (KGD)-peptide (Fig. 2A) (46). The three other ITAs from Aiptasia belong to cnidarian ITA Clade 2, which likewise contains several integrins from Cnidarian species. This clade falls between the PS2 clades and clades containing vertebrate α4/α9 cluster, which have been reported to bind the tripeptide LDV (Fig. 2A) (36). Taken together, the phylogenetic analysis suggests the alpha subunit ITA1 (NM_001434660.1, integrin alpha-6) to potentially bind RGD-containing proteins, while other Aiptasia ITAs may bind to LDV or unknown ligands.

**Figure 2.**
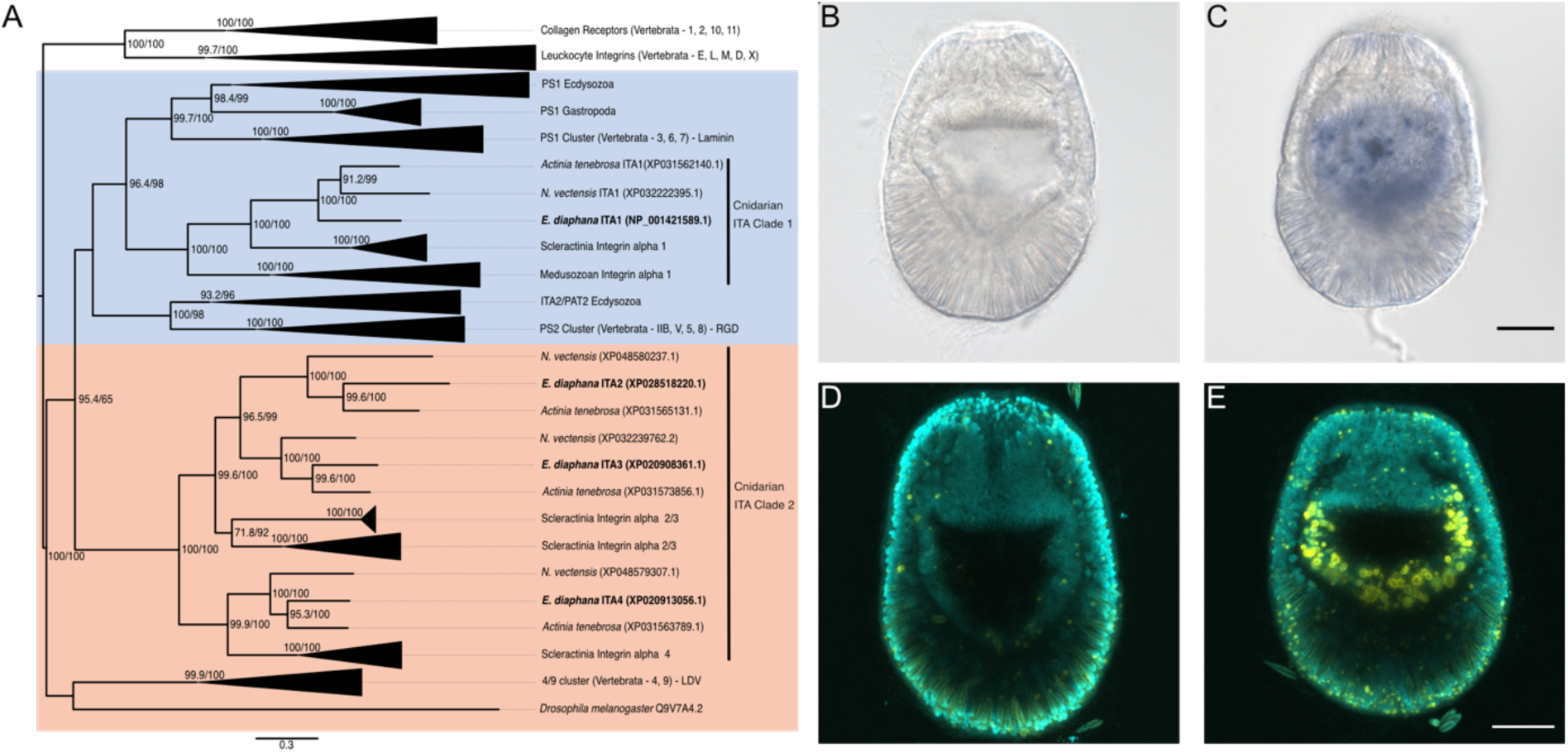
Predicted RGD-binding Aiptasia Integrin alpha 1 is highly expressed in larval endodermal tissue. **A)** Phylogeny based on protein sequences of cnidarian and non-cnidarian integrin alpha sequences. Aiptasia integrin alpha 1 (ITA1) clusters with known RGD and laminin-binding integrins (blue background). The further Aiptasia integrins (ITA2, ITA3, ITA4) cluster in a second group of cnidarian integrins (cnidarian ITA cluster 2) related to LDV binding integrins (peach background). Numbers on Branches and scalebar indicate substitutions per site **B-D)** Localization of ITA1 expression in Aiptasia larvae. ITA1 mRNA was detected with an appropriate antisense probe (C + E) but was absent with a sense probe (B + D) in *in situ* hybridization (B + C) and fluorescent *in situ* hybridization (D + E, yellow signal). Scale bars = 20 µm, cyan (C + E) indicates nuclei.

To determine the location of integrin ITA1 expression we performed *in situ* hybridization (ISH) and fluorescence in situ hybridization (FISH) on Aiptasia larvae. We visualized the expression pattern of the suspected RGD-binding integrin ITA1 (Fig. 2B-E) and found that ITA1 is specifically expressed in the endoderm. Since symbionts are phagocytosed by endodermal cells, this finding is consistent with a possible role of ITA1 in symbiont uptake.

### Integrin blockage inhibits symbiont uptake in symbiotic cnidaria

To assess the role of integrins in symbiont uptake, we exposed aposymbiotic Aiptasia larvae to peptides containing known integrin recognition sequences, LDV (within the peptide EILDV) or RGD (within the peptide GRGDS), and the reverse control peptide SDGRG, to competitively block ligand binding sites before and during symbiont infection (Fig. 3A, B). We found that none of the peptides significantly affected the fraction of infected larvae (data not shown). Additionally, incubation with either EILDV or SDGRG did not significantly affect the number of intracellular symbionts. In contrast, GRGDS-peptide treatment decreased the mean intracellular algae count per larva in a concentration-dependent manner, reducing it to 58% of the uptake seen under SDGRG-peptide-treated or EILDV-peptide-treated conditions (Figure 3B). This suggests that an RGD-integrin interaction plays a crucial part in symbiont uptake.

**Figure 3.**
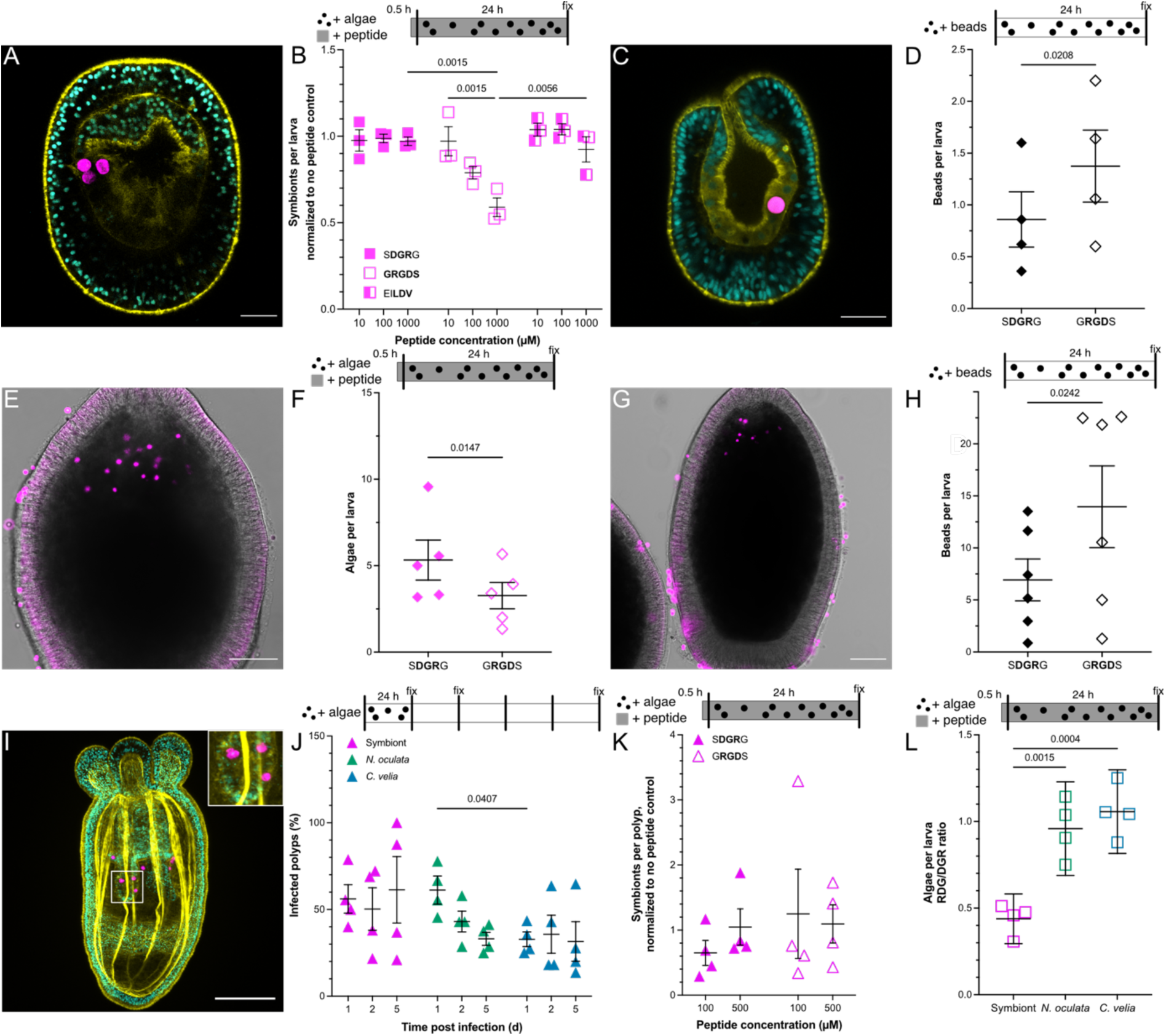
Integrin blockage inhibits symbiont uptake in symbiotic cnidaria. **A)** Aiptasia larvae with symbiont (pink) treated with the control peptide DGR. Cyan = nuclei, yellow = actin. Scale bar = 20 µm. **B)** Symbionts internalized by Aiptasia larvae normalized to a no-peptide control. Aiptasia larvae were exposed to control peptide DGR, integrin ligand RGD or integrin ligand LDV for 30 min then infected with symbionts (*Breviolum minutum*, SSB01) for 24 hours. **C)** Aiptasia larvae with internalized inert bead (pink) Cyan = nuclei, yellow = actin. Scale bar = 20 µm. **D)** Number of peptide-coated beads taken up by Aiptasia larvae. Beads were coated with either control peptide DGR or integrin ligand RGD then incubated with Aiptasia larvae for 24 hours. **E)** *Acropora* larvae infected with SSB01 symbionts (pink) for 24 hours. Scale bar = 100 µm. **F)** Symbionts per *Acropora* larvae after a 24 hour infection and exposure to either the control peptide DGR or integrin ligand RGD (1000 µM peptide). **G)** *Acropora* larvae with internalized inert beads (pink) after 24 hours of exposure. Scale bar = 100 µm. **H)** Control peptide DGR or integrin ligand RGD coated beads per *Acropora* larvae after 24 hours exposure. **I)** Non-symbiotic *Nematostella* polyp with symbionts (pink). Scale bar = 100 µm. **J)** Time course showing the infection rate of *Nematostella* polyps upon symbiont SSB01, or non-symbiont *N. oculata* and *C. velia* exposure. **K)** Symbionts per *Nematostella* polyp after a 24 hour infection and exposure to either the control peptide DGR or integrin ligand RGD. **L)** Algae per larvae as a ratio of RGD- to DGR-treated. Less than 1 indicates RGD decreases algae per larvae compared to the DGR control. For all plots, whiskers depict mean ± SEM. For B, J, K, and L significance was found via ANOVA. For D, F, and H statistical significance found via t-test.

To further investigate this hypothesis, we coated inert polystyrene beads, comparable in size to symbionts (∼8µm diameter) with either GRGDS- or SDGRG-peptides via an oligo-PEG linker which were then incubated with Aiptasia larvae (Fig. 3C, D). After 24 hours, GRGDS-coated beads were phagocytosed by endodermal cells (Fig. S1) and were present inside larvae in higher amounts than beads coated with the control peptide (Fig. 3D). The enhanced uptake or increased early retention of particles coated with RGD-containing peptides indicates that the RGD-integrin interaction is important for phagocytosis.

To explore whether RGD-integrin interactions also facilitate symbiont uptake in corals, we assessed symbiont uptake in *Acropora digitifera* larvae while competitively blocking with either the GRGDS peptide or the control peptide SGDRG (Fig. 3E, F). The number of symbionts in larvae significantly decreased after exposure to 1000µM GRGDS-peptide when compared to the control SDGRG-peptide (Fig. 3F). Further, we found that significantly more beads were taken up intracellularly when coated with GRGDS compared to SDGRG (Fig 3G, H). Taken together, these data suggest that RGD-binding receptors such as integrins promote symbiont uptake and that this interaction is conserved between anemones and corals.

To further characterize the evolutionary conservation of RGD-dependent enhancement of symbiont uptake, we investigated whether the mechanism is also present in the anemone *Nematostella vectensis*. While not naturally symbiotic, *N. vectensis* larvae (tentacle bud stage) can take up low amounts of both symbionts and non-symbiotic microalgae (Fig. 3I, J). To examine the role of the RGD-integrin interaction in the uptake of microalgae in *N. vectensis* larvae we assayed competitive peptide blocking using a range of peptide concentrations (Fig 3K). Notably, the mean symbiont count per larva was not significantly different between GRGDS-peptide treatment and control conditions. These results suggest that the uptake of symbionts is not dependent on recognition by RGD-binding integrins in the non-symbiotic sea anemone *N. vectensis*. Together this indicates that RGD-integrin interactions are utilized by symbiotic Cnidaria to increase symbiont uptake specificity, whereas non-symbiotic Cnidaria rely on more general uptake mechanisms.

As previously mentioned, Aiptasia larvae can phagocytose various microalgae species effectively, both within the Symbiodinaceae family and unrelated non-symbiotic microalgae (43, 44). Hence, we sought to examine whether particle uptake through integrin-dependent phagocytosis was symbiont-specific. To test whether inhibition of integrin binding broadly affects microalgae uptake, we repeated peptide blocking experiments in Aiptasia larvae using symbionts and the non-symbiotic microalgae *Nannochloropsis oculata* and *Chromera velia* (Fig. 3L). We found that symbiont uptake after GRGDS exposure was reduced to less than 50% of the uptake seen with the control peptide (GDGRS) (Fig. 3L). In contrast, we did not observe a significant reduction in the uptake of *N. oculata* or *C. velia* when larvae were treated with the inhibitory GRGDS peptide. These results indicate that integrin-dependent phagocytosis is symbiont-specific.

### Integrin overexpression increases symbiont uptake in HEK293T cells

To further establish that RGD-binding integrins are involved in successful symbiont phagocytosis, we quantified symbiont uptake in human embryonic kidney (HEK) 293T cells that overexpressed the mammalian RGD-binding integrin dimer αVβ3 (38, 45, 46). To assess localization and co-expression of the two subunits, each subunit was tagged with one half of a split-YFP, which led to a robust signal at cell membranes, indicative of correct localization and complex formation (Fig. 4A). We found that overexpression of the mammalian integrins significantly increased the proportion of HEK cells infected with symbionts compared to fGFP-expressing control cells (Fig. 4C, D). Notably, some intracellular symbionts displayed a halo of YFP fluorescence, which indicates the integrin dimers decorated the symbiosome membrane (Fig. 4A - inset). Taken together, symbiont uptake in HEK cells is significantly enhanced by overexpression of RGD-binding integrins.

**Figure 4.**
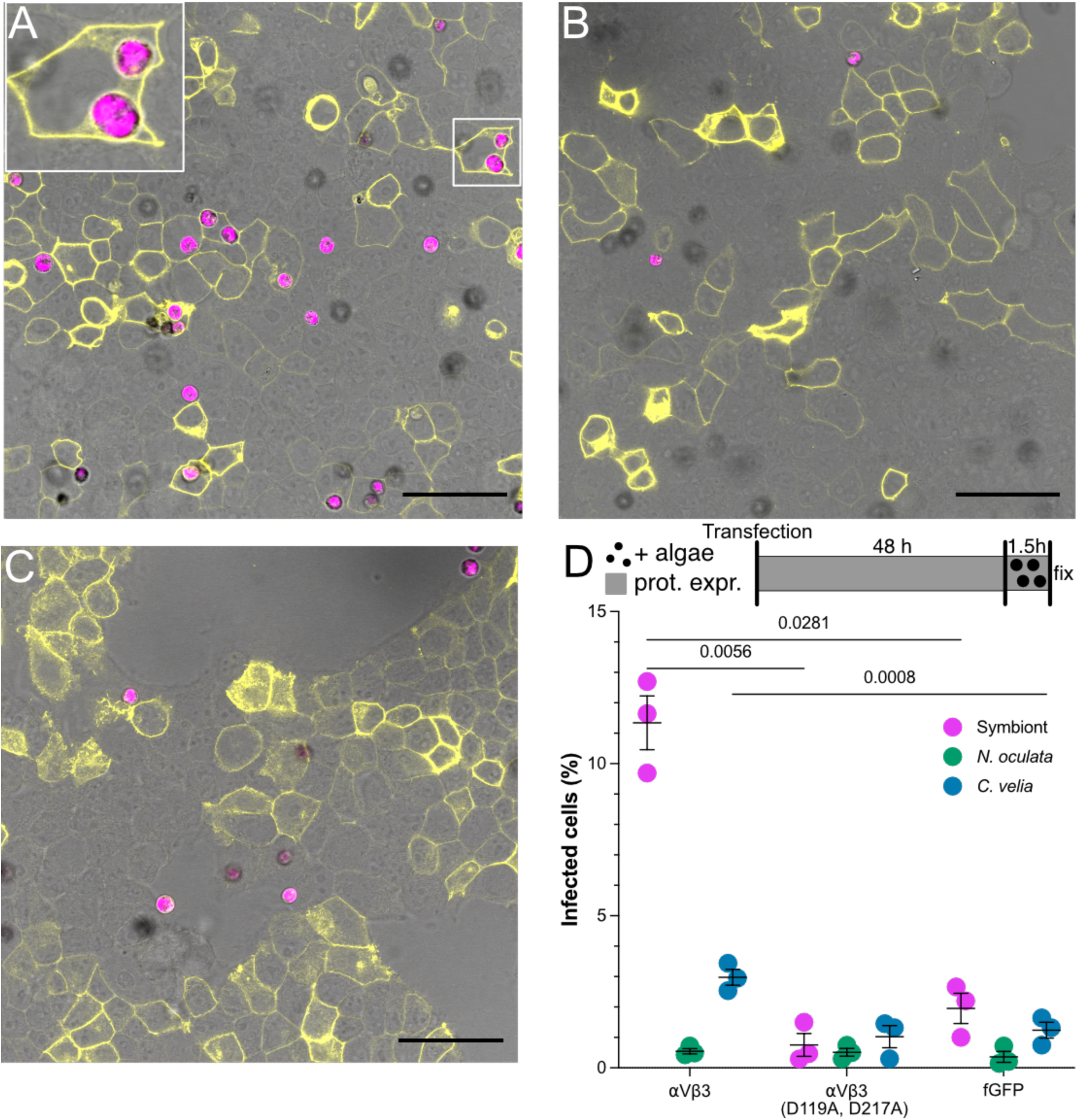
Integrin overexpression increases symbiont uptake in HEK293T cells. **A)** HEK cells transfected with expression plasmids encoding mammalian integrin αV and β3 (each with halves of a split YFP and upon heterodimer formation fluoresce (yellow)) and exposed to symbionts (SSB01, pink). The inset shows integrins localized to the symbiosome membrane. **B)** HEK cells transfected with expression plasmids encoding mammalian αVβ3 integrins with the binding site mutated (D119A, D217A) (each with halves of a split YFP and upon heterodimer formation fluoresce (yellow)) and exposed to symbionts (SSB01, pink). **C)** HEK cells transfected with control plasmid encoding fGFP (yellow) exposed to symbionts (pink). Scale bars for A-C = 50 µm. **D)** Percentage of symbionts (SSB01) and non-symbionts (*N. oculata* and *C. velia*) infected HEK cells that express either αVβ3 integrins, αVβ3 integrins with the binding site mutated (D119A, D217A), or fGFP as a control. Whiskers depict the mean ± SEM. Statistical significance was found via ANOVA followed by Tukey. For all images and plots protein expression was carried out for 48 hours prior to a 90-minute infection with the respective algae.

Integrins bind ligands in a pocket created between the alpha and beta subunit. The conserved cation-binding sites, Metal Ion-dependent Adhesion Site (MIDAS), Adjacent to MIDAS (AMIDAS), and Ligand-associated Metal-binding Site (LIMBS) in the beta-subunit play an important role in binding efficiency (Fig. S2) (47–50). Mutations in aspartic acid residues D119 and D217 (numeration according to processed human integrin beta 3) have been shown to decrease collagen binding, with collagen being the prototypic RGD-containing protein (49). Thus, we generated a double mutant version (D119A, D217A) of the mammalian integrin β3 and used it in the previously described assay (Fig. S2). Overexpression of the integrin β3 mutant, together with integrin αV, showed no difference in the integrin expression pattern (Fig. 4B); however, it resulted in a significantly lower fraction of symbiont-infected HEK cells compared to the wild-type integrin dimer (Fig. 4D). Altogether, this strongly suggests that the recognition of a potential RGD-containing protein on the symbiont surface enhances symbiont uptake, the recognition of which is even conserved in human integrins.

To investigate whether RGD-binding integrins specifically facilitate the uptake of symbionts, we compared uptake of symbionts and non-symbionts (*N. oculata* and *C. velia)* in HEK cells after integrin overexpression (Fig. 4D). We found that symbionts were taken up significantly more frequently than both *N. oculata* and *C. velia* in cells where RGD-binding integrins were overexpressed. Expression of the mutated integrin dimer led to particle uptake similar to the range of uptake in the fGFP-expressing control cells for all algae (Fig. 4D). Although integrin overexpression did significantly increase the number of cells that phagocytosed *C. velia* when compared to fGFP controls, the proportion is still much lower than that of cells containing symbionts upon integrin overexpression (Fig. 4D). Our observations indicate that (1) integrin overexpression enhances symbiont-specific uptake and that (2) mutations in RGD-binding sites negatively impact symbiont uptake which strongly implicates RGD-binding integrins in successful symbiont phagocytosis and symbiosis establishment.

## Discussion

The mechanisms of symbiont uptake in cnidarians are one of the major unresolved questions in the field. Moreover, symbiont uptake is a defining factor when assessing the potential of using heterologous symbionts to improve thermal tolerance in corals. It is unclear whether symbionts are selected through a symbiont-specific uptake route, or if unsuitable algae are expelled following uptake by a more general uptake mechanism. In addition, the specific receptors and respective ligands that mediate uptake are largely unknown (25, 51). Multiple receptor-ligand pairs have been indicated to play a role in symbiont uptake, most noteworthy the glycan-lectin interactions (23, 26, 52). However, sorting post initial uptake, through either apoptosis or vomocytosis has also been reported (43, 53).

To determine additional mechanisms of symbiont selection we used an unbiased transcriptomic approach and identified integrins and their respective downstream signaling molecules to be significantly downregulated specifically in Aiptasia endodermal cells that took up symbionts (54). We hypothesize that integrin signaling is downregulated upon symbiont uptake in a cell intrinsic manner to avoid uptake of multiple symbionts into individual host cells, while this would not necessarily inhibit uptake into neighboring cells. In follow-up experiments, we found that saturation of integrin binding sites with RGD-peptides, a well-known ligand motif of integrins, inhibited symbiont uptake. Specifically, while RGD causes a reduction of the number of symbionts within individual larvae, there is no significant effect on the overall fraction of infected larvae (data not shown). Bay et al. (55) obtained similar results when modifying symbiont surface molecules, and suggested that the observed reduction in uptake was more due to post-phagocytosis retention rather than pre-phagocytosis recognition (55), in line with observed flexibility in the uptake of different Symbiodiniaceae species and even non-symbiotic algae (43, 56). This would indicate that integrin-dependent signaling pathways, which are known to have a multitude of effects on cell motility, cytoskeletal organization, transcription control, proliferation, and cell survival, play an important role in preparing the host cell for a stable symbiosis (35). This is in line with the idea that integrin signaling might be the more ancestral role of integrins rather than cell-adhesion (57).

In addition to their well-known functions in cell adhesion and motility, integrins are also utilized by phagocytic and non-phagocytic cells for pathogen elimination and clearance of apoptotic and tumor cells in flies to humans (38). Integrin activity and binding affinity increases when cytoplasmic proteins interact with the heterodimer to separate their tails and stabilize their high affinity conformation. This is achieved either through inside-out signaling, where alternative signaling modules such as immune signaling alters affinity, or through outside-in signaling, via ligand binding (38). Subsequently, actin remodeling occurs, followed by phagocytosis. Interestingly, many human pathogens, including *Bordetella pertussis*, *Helicobacter pylori*, *Neisseria gonorrhoeae*, *Yersinia pestis*, Adenovirus, Herpes viruses, HIV, SARS-CoV-2 and more, have co-opted integrin receptors to adhere to, enter, enhance colonization and replication, and spread within the host, many using RGD-containing proteins to interact with integrins (42). Integrin-mediated entry is also a method of immune evasion for some pathogens. For instance, professional phagocytes such as macrophages and neutrophils typically phagocytose pathogens via F_c_ receptor-mediated uptake of opsonized pathogens, followed by lysosomal fusion and destruction of the pathogen via oxidative burst, degradative enzymes and toxic peptides. However, some pathogens, such as *Bordetella pertussis*, can bypass this oxidative burst and survive in phagosomes through integrin-mediated entry into phagocytes (58). Pathogens also subvert integrins to invade normally non-phagocytic epithelial and endothelial cells, as well as lymphocytes. Although the exact mechanisms of this phenomenon are not fully understood, the binding of a pathogen by integrins induces integrin clustering and the activation of transduction pathways that initiate internalization (42). These pathogens can also bind to ECM components such as collagen and fibronectin to control cellular signaling and uptake, or degrade ECM components to manipulate integrin functions (42). Given the multitude of pathogens and mechanisms of integrin subversion for entry into host cells, it is not surprising that integrins also play a role in the uptake of endosymbionts. However, to our knowledge, this is the first evidence of such an interaction.

Taken together our findings imply a major involvement of RGD-binding integrins in symbiosis establishment in cnidarians. Several findings, however, leave room for further discussion and investigation. While infection reduction was consistent, blocking with RGD peptides only reduced uptake/retention by ∼50% and had no effect on other algae. This could either imply incomplete/competitive binding of the added RGD peptides, or would be consistent with a complex multistep uptake mechanism. Such an uptake mechanism would likely involve multiple receptors, including glycan-lectin, as well as scavenging receptor interactions and a post-phagocytotic sorting mechanism during early symbiosis establishment (26, 33, 43, 56). In agreement with this, blockage of glycans on the surface of symbionts does not completely inhibit symbiont uptake (21, 25, 26, 29) and, in some cases, has little effect on colonization (31, 55), although these studies all use Aiptasia or coral adult polyps which have unique challenges in quantifying symbiosis. Similar findings have been reported for scavenger receptors and other lectin-like proteins found on the host cell surface, where blockage or RNAi-mediated knockdown leads to inhibition, although incomplete, of symbiont colonization (33, 52). It is likely that glycans, lectins, scavenger receptors and integrins cooperate to distinguish between potential algal partners during uptake and retention. However, the sequence of events and exact partners involved remain to be elucidated.

Additionally, integrin-dependent uptake appears to add specificity, as blocking with RGD peptides only affects symbiont uptake, and does not play a role in the uptake of non-symbiotic algae or in a non-symbiotic host (*N. vectensis*). Similarly, blocking certain glycan/lectin interactions reduces symbiont colonization, but has little effect on heterologous or incompatible symbionts (25). Being able to distinguish between beneficial and potentially parasitic or ineffective partnerships is a critical step in establishing a symbiotic relationship. Our data suggests integrin-ligand interactions to be a potential mechanism to facilitate the uptake of beneficial symbionts. However, non-symbiotic algae are still phagocytosed and establish a transient relationship inside host cells until finally being expelled (43). Therefore, it is unclear whether integrins play a role at the cell surface in the detection of symbionts and facilitation of uptake, or if integrins within the phagocytic vesicle signal to the host whether to expel or incorporate the algae. It is possible that integrins play a role in both steps of symbiosis establishment, uptake and post-phagocytic processing, and is an area of potential future research. In addition, the exact integrin dimer and symbiont cell surface ligand involved in the selection of symbionts is unknown. It is also possible that the host secretes components that bind to symbiont cell wall ligands that are in conjunction recognized by host integrins to facilitate uptake and add an additional layer of specificity into the selection process of beneficial partners, as similar processes have been seen with glycan/lectin interactions in *Xenia* corals (52). Further examination is required to identify integrin ligand pairs, which would be beneficial in the quest to generate optimal host-symbiont pairings.

Integrins are highly conserved molecules that can bind to microbes and coordinate their phagocytosis into multiple cell types across evolutionary scales. Here, we show that RGD-binding integrins of cnidarians facilitate uptake of symbiotic dinoflagellates, thereby identifying them as key players in establishing a healthy symbiosis. Therefore, integrin-dependent phagocytosis may not be specific to pathogen uptake, but instead an evolutionarily ancient mechanism of establishing beneficial intracellular relationships with microbes, that has since been hijacked by certain pathogens. As such, Aiptasia positions itself as a simple yet powerful model organism to uncover the mechanisms of symbiosis establishment and could potentially provide key information for the generation of symbiont host pairings resistant to the challenges faced by coral reefs in today’s changing climate.

## Materials and Methods

### Live organism and cell culture

#### Microalgae maintenance

*Breviolum sp.* clade B (family Symbiodiniaceae, strain SSB01; symbiont) (59), *Microchloropsis gaditana* CCMP526 (National Center for Marine Algae and Microbiota, Bigelow Laboratory for Ocean Sciences), *Nannochloropsis oculata*, and *Chromera velia* (Norwegian Culture Collection of Algae K-1276; Norwegian Institute for Water Research) were grown in cell culture flasks in 0.22 μm filter-sterilized Diago’s IMK medium (Wako Pure Chemical Corporation). *Breviolum sp., N. oculata* and *C. velia* were cultured at 26 °C, whereas *M. gaditana* was maintained at 18 °C. Prior to infection, all microalgae cultures (including *M. gaditana*) were kept at 26 °C on a 12 hr light / 12 hr dark cycle under ∼ 20 - 25 μmol m^-2^s^-1^ of photosynthetically active radiation (PAR), as measured with an Apogee PAR Quantum meter (MQ-200; Apogee), 1 - 2 weeks post-splitting.

#### Aiptasia stock culture conditions

Clonal Aiptasia (*Exaiptasia diaphana*) lines F003 and CC7 (Carolina Biological Supply Company; 162865) were maintained in translucent polycarbonate tanks (GN 1/4 - 100 cm (# 44 CW) and 1/9 - 65 cm (#92 CW); Cambro, Huntington Beach, CA, USA) filled with artificial seawater (ASW; PRO-REEF Sea Salt, Tropic Marin^®^) at 31 – 34 ‰ salinity. Aiptasia stocks were kept in Intellus Ultra Controller Incubators (I-36LL4LX; Percival, Perry, USA) at 26 °C on a diurnal 12 hr light:12 hr dark cycle (12L:12D) under white fluorescent bulbs with an intensity of ∼ 20 - 25 μmol m^−2^s^−1^ of PAR. Animals were fed twice per week using freshly hatched *Artemia* nauplii (Ocean Nutrition^TM^) and cleaned 3 hr later with cotton-tip swabs and tissue paper, followed by a water change.

#### Aiptasia spawning and larval culture conditions

Spawning of Aiptasia clonal lines F003 and CC7 was induced as described previously (60). Developing Aiptasia larvae were maintained in glass beakers in filter-sterilized artificial seawater (FASW) at 26 °C and exposed to a 12L:12D cycle.

#### *Nematostella* stock culture conditions

*Nematostella vectensis* stocks were cultured in polycarbonate tanks filled with 1/3 ASW at 11.0 – 11.5 ‰ salinity. Tanks were kept in darkness at 18 °C, and animals were fed once to twice per week with freshly hatched *Artemia* nauplii. *N. vectensis* were transferred into clean tanks filled with fresh 1/3 ASW (18 °C) once a week.

#### *N. vectensis* spawning and larval culture conditions

Spawning was induced as previously described (39, 61), with the following adaptations: Female and male tanks were rotated to spawn every 2 - 3 weeks. The day before spawning, animals were transferred to clean tanks with fresh 1/3 ASW (18 °C) and incubated for 8 hr at 26 °C under white fluorescent bulbs with an intensity of ∼20 - 25 μmol m^-2^s^-1^. The water was replaced with fresh 1/3 ASW (18 °C) the next day and tanks were monitored for spawning for 2 – 3 hr. Egg packages were transferred into petri dishes and fertilized with sperm water. Developing larvae were kept at 18 °C or 26 °C and filtered into fresh 1/3 FASW after escaping the jelly coat.

#### Acropora digitifera spawning

Colonies of the coral *Acropora digitifera* were collected off Sesoko Island (26 3°7’41”N, 127 5°1’38”E, Okinawa, Japan) according to Okinawa Prefecture permits and CITES export and import permits. They were handled as previously described (44) at Sesoko Tropical Biosphere Research Center (University of Ryukyus, Okinawa, Japan). Isolated *Acropora* colonies were kept until spawning, and spawned symbiont-free gametes were mixed for fertilization. Planula larvae were then maintained at around 1000 larvae/L in filtered natural seawater, which was exchanged daily.

#### HEK293T cell culture conditions

Adherent HEK293T cells (American Type Culture Collection, VA, USA) were cultured in Gibco^TM^ Dulbecco’s Modified Eagle’s Medium (DMEM; 41966-029; Thermo Fisher Scientific^TM^) supplemented with 10 % (vol/vol) heat-inactivated Gibco^TM^ Fetal Bovine Serum (FBS; 10500-064; Thermo Fisher Scientific^TM^) and 1 % (vol/vol) Gibco^TM^ 10,000 U/mL penicillin-streptomycin (15140-122; Thermo Fisher Scientific^TM^). Cells were grown in cell culture flasks (658195, Cellstar^®^; Greiner Bio-One) at 37 °C with 5 % CO_2_ in a HERAcell^TM^ 150i cell incubator (50116047; Thermo Fisher Scientific^TM^) and passaged two to three times a week using 0.25 % Gibco^TM^ Trypsin-EDTA (25200-056; Thermo Fisher Scientific^TM^).

### Infection assays

#### Bead Coating

COMPEL™ Magnetic beads with COOH modification (excitation 480 nm, emission 520 nm, Bangs Laboratories, UMDG003) were coated with peptides and used for infections. Around 11 million Beads were washed 3x in 400 µL MES buffer (0.05 M, pH 5, Sigma-Aldrich, M8250) and resuspended in 320 µL MES buffer. Next, 40 µL each of ED(A)C (50 mg/mL, *N*-(3-Dimethylaminopropyl)-*N*′-ethylcarbodiimid -hydrochlorid, Sigma Aldrich, E6383) and Sulfo-NHS (50 mg/mL, sulfo-N-hydroxysuccinimide, Abcam, ab14569) were added and incubated with rotation at room temperature (RT) for 15 min, after which they were washed 3x in 400 µL ice-cold MES buffer and resuspended in 390 µL ice-cold phosphate buffered saline (PBS, pH 7.2). A linker was then added to the beads through the incubation with the addition of 10 µL of 100 mM NH2-PEG8-Propionic acid (Sigma-Aldrich, JKA12005) for 2 hr at RT with rotation. After this, the beads were washed and incubated in 400 µL Tris (50 mM, pH 7.5, Carl Roth, 4855.2) for 15 min, and then washed 3x in 400 µL MES buffer. At this stage beads can be stored before coupling to peptides. 0.5 million beads were used for coupling and resuspended in 160 µL MES buffer. 20 µL each of ED(A)C and Sulfo-NHS (50 mg/ml each) were added and incubated at RT with rotation for 15 min, followed by 3 washes in MES buffer. Beads were resuspended in 99 µL PBS (pH 7.2) and 1 µL 100 mM of peptide, SDGRG (H-Ser-Asp-Gly-Arg-Gly-OH, Bachem, 4015321), GRGDS (H-Gly-Arg-Gly-Asp-Ser-OH, Bachem, 4008998) or EILV (FibronectinCS-1 Fragment, Bachem, 4026203) was added and incubated with rotation at RT for 2 hr. Beads were then incubated at RT with rotation for 15 min in Tris (50 mM, pH7.5), before 3 final washes in MES buffer and resuspension in FASW.

#### Infection of Aiptasia larvae, *A. digitifera* larvae and *N. vectensis* tentacle bud stages

At least three replicates of Aiptasia larvae (4 - 8 dpf), *A. digitifera* larvae (3-6 dpf) or *N. vectensis* early tentacle bud stages (4 - 6 dpf) were collected in 1.5 mL bovine serum albumin coated (BSA; A7906; Sigma-Aldrich) tubes. Where indicated 500 μL SDGRG, GRGDS, or EILDV peptide (stocks: 100 mM) solutions were prepared in FASW (Aiptasia) or 1/3 FASW (*N. vectensis*) at indicated concentrations. 50 *N. vectensis* tentacle buds or 300-500 Aiptasia larvae in 500 µL 1/3 FASW or FASW, respectively, were transferred to each peptide solution using 1 % BSA-coated pipette tips and incubated for 30 min at 26°C, when treated with peptides, otherwise this step was omitted. Infection with algae of beads were performed at a final concentration of 5 x 10^4^ cells/mL (Aiptasia/*A. digitifera*) or 1 × 10^5^ cells/mL (*N. vectensis*) and incubated at 26 °C with rotation (1 rpm) and exposed to a 12L:12D cycle for 24 h. Prior to fixation, polyps were relaxed in 7 % magnesium chloride hexahydrate (MgCl_2_-6 H_2_O; LC-5041.4; Labochem^®^ International) in 1/3 FASW for 10 min and transferred into 1 % BSA-coated tubes.

#### Staining and mounting

Samples were fixed in 4 % formaldehyde solution (F1635; Sigma-Aldrich) for 30 min and then washed twice in 0.1% Triton X-100 in phosphate-buffered saline (PBS-Triton) (3051; Carl Roth). They were either stained for f-actin and DNA (see below) or directly washed stepwise into glycerol from 30 % to 50 % and finally mounted in 87 % glycerol (G5516; Sigma-Aldrich) in PBS with the addition of 2.5 mg/mL 1,4-diazabicyclo[2.2.2]octane (DABCO; D27802; Sigma-Aldrich). A lash sword was used to position the polyps along their lateral axis on the microscopy slide. Non-toxic double-sided tape (TES5338; tesa^®^) was used as a spacer between the microscopy slide and the coverslip. For staining, fixed Aiptasia larvae or *N. vectensis* polyps were washed 3 times in 0.05 % Tween20 (P7949; Sigma-Aldrich) in PBS (PBS-T) for 5 min. For permeabilization, samples were rotated at 0.25 rpm in 1 % PBS-Triton and 20 % dimethyl sulfoxide (DMSO; 67-68-5; Thermo Fisher Scientific^TM^) for 1 hr at RT. After 3 washes in PBS-T for 10 min, larvae or polyps were incubated in Phalloidin-Atto 565 (94072; Sigma-Aldrich) diluted 1:200 in 0.05 % PBS-T for 1 hr at 0.25 rpm in the dark. Samples were washed 3 times in 0.05 % PBS-T for 5 min before incubation with 10 μg/mL Hoechst (B2883; Sigma-Aldrich) diluted in Tris-buffered saline (pH 7.4), 0.1 % Triton X-100, 2 % BSA, and 0.1 % sodium azide (S2002; Sigma-Aldrich) for 20 - 30 min at 0.25 rpm and RT in the dark. Larvae or polyps were washed 3 times for 5 min with 0.05 % PBS-T and then mounted as described above.

#### Microscopic analysis

Confocal microscopic analysis was carried out on a Leica TCS SP8 confocal laser scanning microscope using a 10X dry immersion objective (numerical aperture = 0.30) or a 63X glycerol immersion objective (numerical aperture = 1.30), Leica LAS X and Fiji software (version 2.1.0/1.53c) (62). Hoechst, Atto-565, and microalgae autofluorescence were excited with 405, 561 and 633 nm laser lines, respectively. Fluorescence emission was detected at 410-501 nm for Hoechst, 542-641 nm for Phalloidin-Atto 565, and 645-741 nm for symbiont autofluorescence.

#### Quantification of infection efficiency

The number of intracellular algae or particles was counted for at least 30 Aiptasia larvae, 20 *A. digitifera* larvae or 30 *N. vectensis* polyps per replicate per microalgal or particle type and data recording was documented in Microsoft Excel version 16.16.6.

### Molecular cloning

#### RNA isolation/cDNA synthesis

1-2 CC7 polyps or 5000 aposymbiotic Aiptasia larvae (5 – 8 dpf) were homogenized until dissolved using 1 mL TRIzol^®^ (15596018; ambion^®^ by life technologies; Thermo Fisher Scientific^TM^)) and a tissue homogenizer (MiniBatch D-1; MICCRA). After samples were incubated at RT for 5 min, 200 µL Chloroform (132950; Honeywell) was added, and the samples were incubated for 3 min at RT before centrifugation (10,000 - 12,000 × g, 15 min, 4 °C). The supernatant was transferred into a fresh RNase-free 1.5 mL Eppendorf tube and one volume of 70 % ethanol was added. RNA was applied to RNeasy Mini Kit columns (Qiagen, 74104) and purified according to manufacturer’s instructions. cDNA was transcribed using the SuperScript^®^ IV Reverse Transcriptase (InvitrogenTM, 18090010; ThermoFisher Scientific^TM^) following manufacturer’s instructions. Complementary RNA was removed by adding 0.5 μL of 5 U/μL *E. coli* RNase H (MO2975; New England Biolabs Inc. (NEB)) followed by incubation for 20 min at 37 °C.

#### Restriction cloning

For cloning of plasmid P-0251, the insert was PCR-amplified from cDNA using Q5^®^ High-Fidelity DNA Polymerase (M0491; NEB) with the primers and template defined in Table 2. After column- or gel-purification using the GeneJET PCR Purification Kit (Thermo Fisher Scientific^TM^) according to manufacturer’s manual, PCR product and vector (Table 1+2) were digested using enzymes indicated in Table 2, according to manufacturer’s instructions (New England Biolabs), ligated using T4 DNA ligase (NEB) according to manufacturer’s instructions and transformed to TOP10 chemically competent *E. coli* (Thermo Fisher Scientific^TM^) according to manufacturer’s instructions. The insert sequence was checked by sequencing.

**Table 1.** Plasmids used in (F)ISH and cell-culture experiments.

| **INTERNAL**<br>**NUMBER** | **INSERT** | **BACKBONE** | **FLUOROPHORE** | **ADDGENE**<br>**NUMBER** |
| --- | --- | --- | --- | --- |
| **P-0062** |  | pCRII-TOPO/8c + ASCI&PacI restriction sites |  |  |
| **P-0251** | Aiptasia Probe ITA1 | pCRII-TOPO/8c + ASCI&PacI restriction sites |  |  |
| **P-0266** | Human Alpha-V-Integrin | mEmerald-N1 | mEmerald | 53985 |
| **P-0268** | Mouse Integrin-Beta3 | mEmerald-N1 | mEmerald | 54130 |
| **P-0273** | farnesylated eGFP | pEGFP (clonotech) | eGFP |  |
| **P-0299** | Human Alpha-V-Integrin |  | C-term-split-YFP |  |
| **P-0300** | Integrin-Beta3 |  | N-term-split-YFP |  |
| **P-0301** | Empty | pCS2+ |  |  |
| **P-0302** | Integrin-Beta3 (D119A) |  | N-term-split-YFP |  |
| **P-0303** | Integrin-Beta3<br>(D119A+D217A) |  | N-term-split-YFP |  |

**Table 2.**
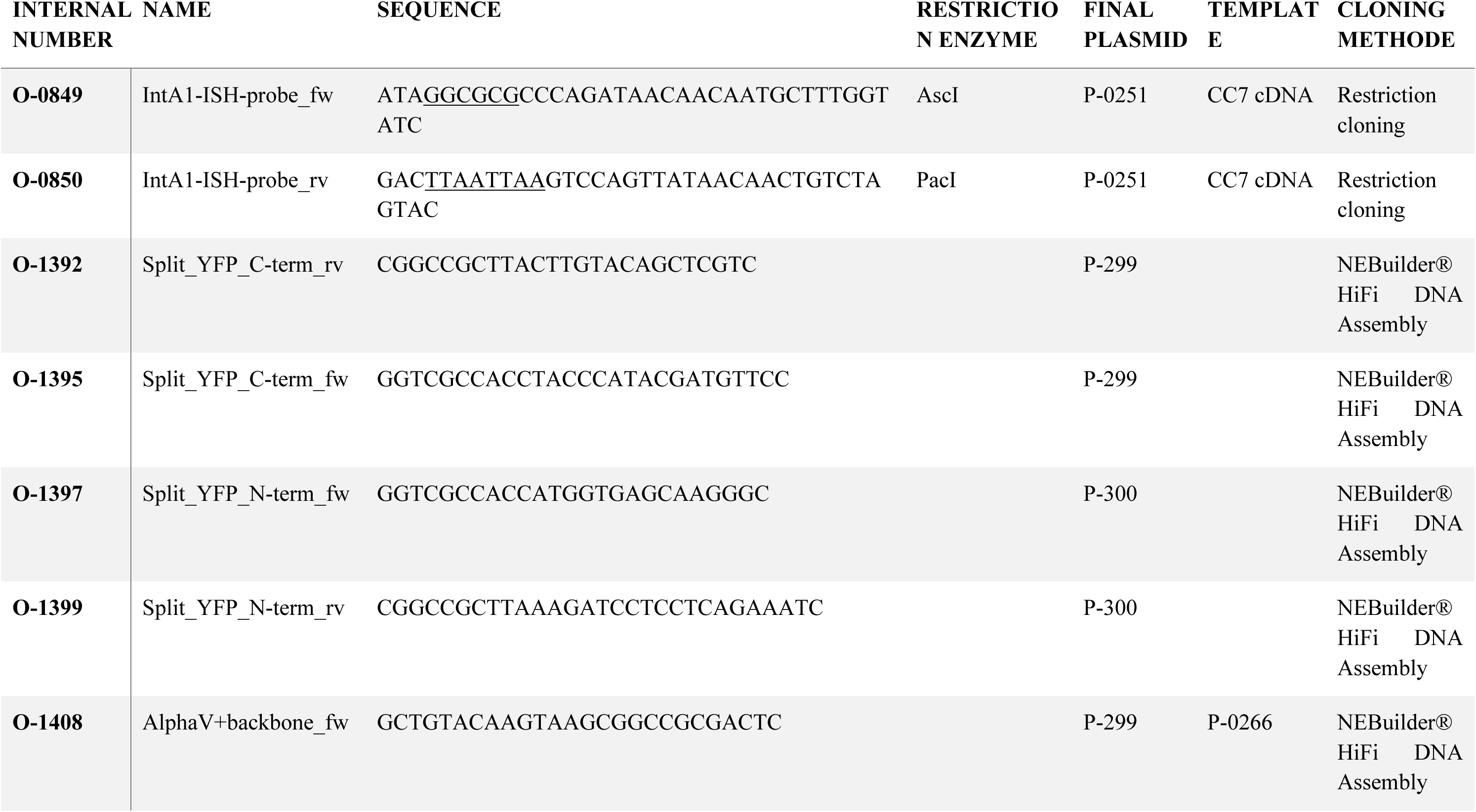
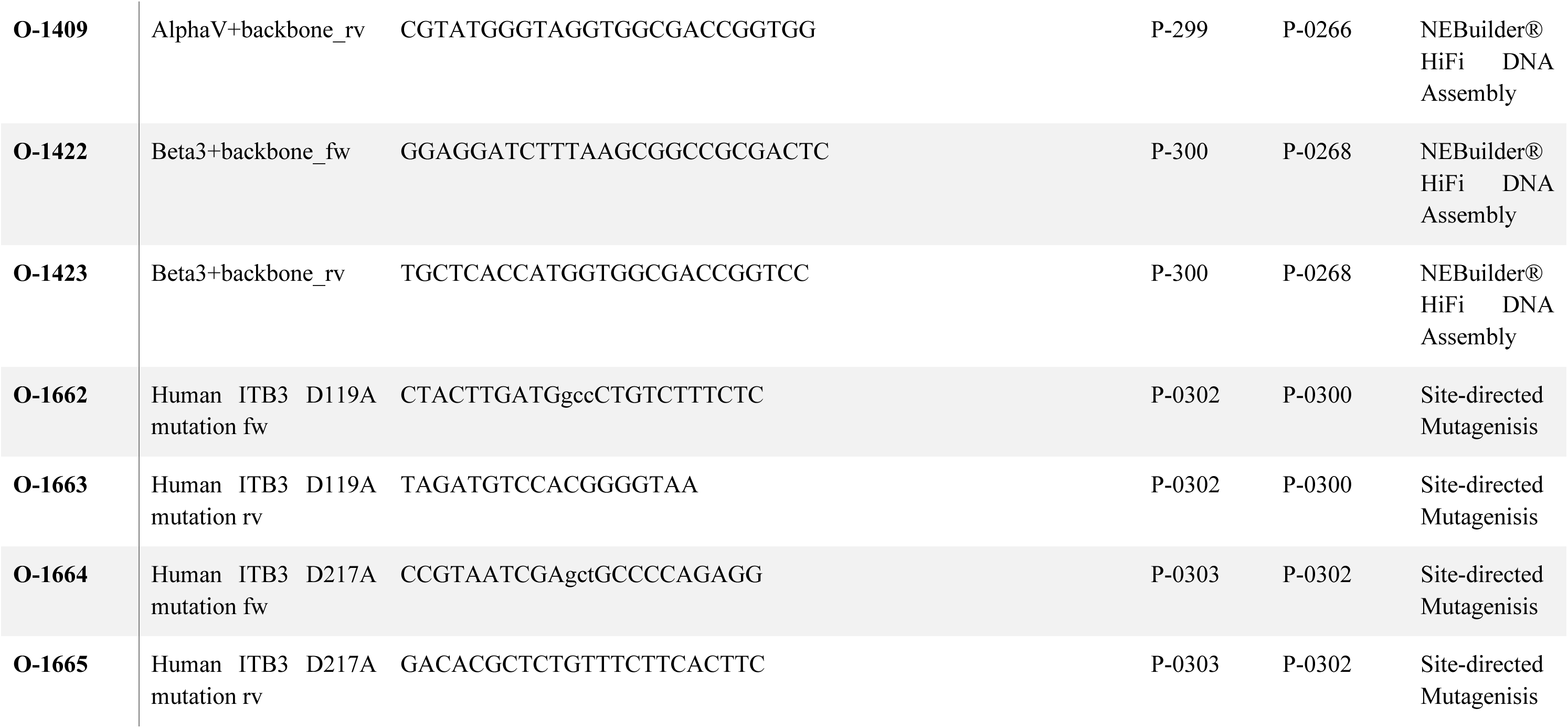
Primers used for construction of plasmids.

#### NEBuilder® HiFi DNA Assembly

For cloning of plasmids P-0299 and P-0300, fragments were amplified with primers from cDNA or plasmid templates as indicated in Table 2, using Q5^®^ High-Fidelity DNA Polymerase, per manufacturer’s instructions. After column- or gel-purification of inserts using the GeneJET PCR Purification Kit (Thermo Fisher Scientific^TM^) according to the manufacturer’s manual, cloning vectors (Table 1) were digested using enzymes indicated in Table 2, according to manufacturer’s instructions (NEB). Template plasmids were removed by gel extraction using the GeneJET PCR Purification Kit. Vector and insert concentrations were determined by 1 % agarose gel electrophoresis, and constructs were assembled using the NEBuilder^®^ HiFi DNA Assembly Cloning Kit (E5520S; NEB) using a 1:2 vector:insert dsDNA pmols ratio following manufacturer’s instructions. Samples were diluted 1:4 - 1:3 prior to transformation of 2.5 μL into chemically competent DH5-alpha *E. coli* (18265017; Thermo Fisher Scientific^TM^) or Endura^TM^ DUOs bacteria (60240-0; Lucigen; BiosearchTM Technologies). The insert sequence was checked by sequencing.

#### Site-directed mutagenesis

For cloning of plasmids P-302 and P-0303, primers with mismatches (underlined in Table 2) were used to amplify the original plasmid (Table 1) using the Q5^®^ High-Fidelity DNA Polymerase, per manufacturer’s instructions. Similar to site-directed mutagenesis kits, 1 µL of PCR product was used in a reaction mix containing, 1 µL of T4 DNA ligase buffer (B0202S; NEB), 1 µL T4 Polynucleotide Kinase (M0201S; NEB), 1 µL T4 ligase (M0202S; NEB), 1 µL DpnI (R0176L; NEB) and 5 µL H_2_O, and incubated for 1 hr at 37 °C. Half the reaction mix was transformed into chemically competent DH5-alpha *E. coli*.

### HEK293T cell infection assays

#### Mammalian αVβ3 plasmid constructs

If not stated otherwise, 750 ng human integrin subunit αV fused to the C-terminal part of a split-YFP (P-0299), and 750 ng mouse integrin subunit β3(P-0300)) fused to the N-terminal part of a split-YFP were used to overexpress the integrin dimer αVβ3 in human HEK293T cells. 250 ng pEGFP-f (farnesylated eGFP, AGP57; Clontech; obtained from Ary Shalizi, Stanford University, P-0273) together with (empty) 750 ng pCS2+ plasmid constructs (P-0301, kind gift from Sergio Acebron, Heidelberg University) were transfected as a negative control. As non-functional control the beta 3 subunit was mutated by changing aspartic acid D119 and D217 to alanine (P-0303, based on (49)), and overexpressed together with the alpha subunit as indicated above.

#### Calcium phosphate transfection

HEK293T cells (0.75 x 10^5^ cells/well) in DMEM were grown in 12-well plates (665102; Cellstar^®^; Greiner Bio-One) on sterile Poly-L-lysin (0.01%, P8920; Sigma Aldrich) coated coverslips overnight at 37 °C with 5 % CO_2_. Appropriate amounts of plasmid DNA constructs (see above) and 7.5 μL 2.5 M CaCl_2_, were added to water to a final volume of 75 µL. An equal volume of 2X HeBS buffer (pH 7.05) was added dropwise, and the mix was incubated for 10 min at RT. The transfection mix was added and cells were incubated at 37 °C with 5 % CO_2_ for 5 - 7 hr before gently washing twice with 1X dPBS and subsequently adding fresh DMEM. Cells were incubated for 48 hr prior to infection.

#### Infection of HEK293T cells

HEK293T cells were infected using symbionts, *N. oculata*, or *C. velia* 48 hr post-transfection. 1 - 2-week-old algae cultures in IMK medium were pelleted for 5 min at 2000 g and resuspended in DMEM at a final concentration of 3 × 10^5^ algae cells/mL (if not stated otherwise). Prior to infection, transfected HEK293T cells were washed once with pre-warmed 1X dPBS, before adding 1 mL of algae suspension in each well. After incubation for 1.5 hr or 4 hr (as indicated) at 37 °C with 5 % CO_2_, cells were washed once with pre-warmed 1X dPBS. HEK293T cells were fixed using 4 % PFA in 1X dPBS at RT for 30 min, washed with dPBS and mounted on microscopy slides in 100 % glycerol. Algal uptake was compared in a minimum of three independent biological replicates.

#### Quantification of HEK293T cell infection efficiency

The total number of transfected HEK293T cells and transfected cells with intracellular algae was counted in Z-stacks (size 20 - 40 μm) of six randomly chosen fields of view per sample per microalgal type. Confocal microscopic analysis was carried out on a Leica TCS SP8 confocal laser scanning microscope using a 63X glycerol immersion objective (numerical aperture = 1.30), Leica LAS X and Fiji software (version 2.1.0/1.53c) (62). eGFP, YFP, and algae autofluorescence were excited using 488, 514, and 633 nm laser lines, respectively. Fluorescence emission was detected at 493 - 570 nm for eGFP, 519 - 590 nm for YFP, and 645 - 741 for microalgae autofluorescence.

### (Fluorescent) *In situ* hybridization

#### Probe generation

The plasmid for probe-generation (Table 2) was designed as described in the molecular cloning section. Digoxygenin-labeled probes were generated after linearization with either AscI or PacI (NEB; dependent on final orientation of the probe) using Roche DIG RNA Labeling Kit (SP6/T7). In short, 1 µg linearized plasmid was incubated with 10 x transcription buffer (1X final conc.), 10x NTP labeling mixture (1X final conc.), RNase inhibitor (1 U/µL final conc.), T7 or Sp6 Polymerase (1 U/µL final conc.) for 2 hr at 37 °C, before adding DNaseI (1 U/µL final conc) for an additional incubation of 20 min. RNA was extracted using phenol/chloroform/isoamyl alcohol (25:24:1, Carl Roth), and precipitated using 3 M sodium acetate (Sigma-Aldrich) and isopropanol (Sigma-Aldrich).

#### *In situ* hybridization

For *in situ* hybridization Aiptasia larvae (3-8 dpf) were fixed in 3.7 % PFA in FASW for 1 hr, washed 3 times in PBS-Triton + 1 % BSA, followed by 3 washes in 100 % methanol (Sigma-Aldrich). Larvae were then incubated in 90 % methanol (Sigma Aldrich) and 3 % hydrogen peroxide in water for 30 min at RT, followed by rehydration through 60 % methanol and 40 % of 0.1 % Tween20 in PBS (PBT), 30 % methanol and 70 % PBT and 4 washes in 100 % PBT for 5 min each. Next, larvae were incubated in 0.01 mg/mL proteinase K (Roche) in PBT for 8 min prior to two 3 min washes in 2 mg/mL glycine in PBT. Larvae were washed with 1 % Triethanolamine (Merck) in PBT, then with 0.3 µL/mL acetic anhydride in 1 % triethanolamine in PBT for 10 sec, then with 0.6 µL/mL acetic anhydride (Grüssing) in 1 % triethanolamine in PBT for 10 sec and then twice in PBT, followed by a 30 min incubation in 4 % PFA in PBT at RT and 5 washes in PBT.

Next, larvae were incubated for 10 min in hyb buffer (50% formamide (Sigma Aldrich), 0.075 M trisodium citrate (Gerbu), 0.75 M sodium chloride (Sigma Aldrich), 0.05 mg/mL heperin (Serva), 0.25 % Tween-20, 1 % SDS, 0.05 mg/mL salmon sperm DNA (Invitrogen), pH 4.5) at RT, followed by 1 – 4 hr incubation in fresh hyb buffer at 60 °C. Larvae were then incubated with denatured (10 min at 90 °C) DIG-labeled probes (1 ng/µL) at 60 °C for 36 – 72 hr with rotation. For competition with unlabeled probes a 10-fold excess was used. After incubation, larvae were washed at 60 °C for 5 min in hyb buffer, and again for 15 min in hyb buffer, prior to wash into 2X SSC buffer (0.3 M sodium chloride, 0.03 M trisodium citrate, pH 7.0) in consecutive steps of 75 % hyb buffer + 25 % 2X SSC buffer, followed by 50 % each buffer and 25 % hyb buffer + 75 % 2X SSC buffer and finally 100 % 2X SSC buffer for 10 min each at 60 °C followed by two 20 min washes at 60 °C in 0.05X SSC (7.5 mM sodium chloride, 0.75 mM trisodium citrate, pH 7.0). Next, larvae were washed into PBT in three steps at RT from 75 % 0.05X SSC + 25 % PBT to 50 % each, 25 % 0.05X SSC + 75 % PBT to 100 % PBT. After this, larvae were incubated in blocking buffer (Blocking solution, Roche in Maleic acid buffer) for 30 min at RT, followed by incubation in anti-Dig alkaline phosphates (for *in situ* hybridization) or anti-Dig horseradish peroxidase (for fluorescent *in situ* hybridization) diluted 1:5000 in blocking buffer at 4 °C overnight, followed by 3 short washes in PBT and 7 washes of 20 – 30 min at RT.

For detection of *in situ* hybridization, larvae were washed twice in AP buffer (0.1 M sodium chloride, 0.1 M tris (pH 9.5), 0.1 % Tween-20) for 5 min at RT and twice in AP buffer + 0.05 M magnesium chloride for 5 min at RT. Larvae were then incubated in NBT/BCIP staining solution (0.1 M trisHCL (pH 9,5), 0.1 M sodium chloride, 1:50 NBT/BCIP stock (Roche)) at 37 °C until sufficient staining was detected, at which point staining was stopped with an equal volume of 100 % ethanol. This was followed by 2 washes in 100 % ethanol and 3 washes in PBS-Triton followed by a final wash with PBS, before mounting in 90 % glycerol, 10 % Tris (0.1 M, pH8.0).

For detection of fluorescent *in situ* hybridization, the TSA Plus kit (Perkin Elmer) was used. Larvae were incubated for 10 min in FITC stock solution diluted 1:50 in amplification diluent at RT, after which they were washed twice in PBS-Triton for 5 min at RT. For counterstaining of the nuclei 10 μg/mL Hoechst was used for 10 min at RT, followed by three 10 min washes at RT in PBS-Triton and one wash in PBS. The samples were then mounted in 90 % glycerol, 10 % Tris (0.1 M, pH 8.0) + 2.6 mg/mL DABCO, and microscopically analyzed as described above.

### RNA sequencing and analysis

Data from Jacobovitz, *et al.* (43) were used and analysed as described within. Shortly, Aiptasia larvae (300/mL) were infected with SSB01 or *M. gaditana* (10^5^ cells/mL) or left uninfected for 24 hours. 3-5 larvae per replicate were incubated in calcium magnesium free seawater (CMF-ASW) for 5 minutes then incubated for 2min 0.5% Pronase (Sigma; 10165921001) and 1% sodium thioglycolate (Sigma; T0632) diluted in CMF-ASW and pipetted up and down to detach the endodermal layer. The endoderm was transferred to FASW and pools of 7-20 cells for each replicate were picked using a microcapillary needle (Science Products; GB100T-8P). Capillaries contained 4.3 µL lysis buffer (0.2% Triton X-100, 1 U µL −1 Protector RNase Inhibitor (3335399001; Sigma–Aldrich), 1.25 µM oligo-dT30VN and 2.5 mM dNTP mix) and cells were flushed out of the capillary with lysis buffer, then flash frozen. Sequencing libraries were prepared by RNA reverse transcription and pre-amplification of complementary DNA before being sequenced on NextSeq 500 (Illumina) with 75-base pair paired-end reads.Raw sequences deposited at National Center for Biotechnology Information Sequence Read Archive (SRA) (see ‘Data availability’ statement). Genes of interest for the analysis were manually selected from the integrin signaling pathways as described in (54).

### Integrin alpha (ITA) phylogeny

Identification of integrin ⍺ proteins was performed in Geneious 10.2.6 (https://www.geneious.com). Aiptasia integrin ⍺ genes with refseq accession numbers XP_0285182201.1, XP_028518220.1, XP_020908361.1, and XP_020913056.1 (we propose referring to these as ITA1, ITA2, ITA3, and ITA4 respectively) were used as starting point for a blastp search (in Geneious) against the refseq protein database, retrieving the 40 top hits for each, combining these to a total of 83 unique hits. Several of these were different isoforms which were manually curated to give a total of 49 unique hits. The same proteins together with the human ITAE protein (Uniprot accession P38570) were used for a blastp search against the uniprot database retrieving 30 top hits each, combined to a total of 55 unique hits. Sequences were aligned using the MUSCLE Alignment function with default parameters (in Geneious). The protein alignment was then imported into IQ-TREE (1.6.12) and used to perform phylogenetic analysis (63). First, ModelFinder determined that the best mode of protein evolution was WAG+I+G4 model (64), using the Bayesian information criterion. Phylogenetic trees were then generated using 1,000 ultrafast bootstrap iterations (65) and the SH-aLRT test (66). The tree was visualized using FigTree (http://tree.bio.ed.ac.uk/software/figtree/). The full tree (Fig. S3) was then collapsed for ease of analysis.

### Data availability

Raw reads of the RNA sequencing data can be accessed at the National Center for Biotechnology Information SRA with the following accession numbers: SRX7119772–7119776 (cells from aposymbiotic larvae), SRX7119782–7119787 (symbiotic cells) and SRX7119777–7119781 (aposymbiotic cells from symbiotic larvae) (combined in the SRA project SRP229372); and SRX7229078–7229080 (*M. gaditana*-containing cells) and SRX7229075–7229077 (microalgae-free cells from *M. gaditana*-containing larvae) (combined in the SRA project SRP233508).

### Statistics

Data was analyzed using Prism 9 (Version 9.1.1; GraphPad Software, LLC).

Descriptive statistics (mean and standard deviation) and SEM were calculated. When two parameters were compared, paired parametric t-tests were used to calculate p-values (significance level p < 0.05). When more than two parameters were analyzed, 2-way ANOVA was fitted on the data, and Šídák’s or Tukey’s multiple comparison tests were performed to determine statistical significance for each comparison (alpha = 0.05).

## Acknowledgements and funding sources

### Acknowledgements

We would like to acknowledge feedback and editing support from Sami El Hilali and Benjamin Jenkins.

## Author contributions

Conceptualization, V.A.S.J, S.R., and A.G.; methodology, V.A.S.J, M.D., I.S., S.R., S.G.G., and I.M.; software, S.G.G. and P.A.V.; formal analysis, V.A.S.J, M.D., S.R., J.S., and S.G.G.; data interpretation, V.A.S.J, M.D., S.R. and A.G.; resources, A.G.; writing—original draft, M.D., S.R. and M.F.; writing— review and editing, S.R., M.F., M.D., and A.G.; project administration; V.A.S.J, S.R., and A.G.; funding acquisition, A.G.

## Supplementary figures

**Figure S1.**
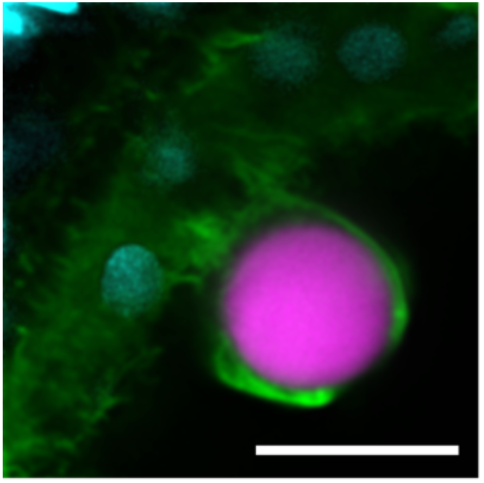
RGD-coated beads are phagocytosed by Aiptasia endodermal cells. Aiptasia larvae exposed to inert beads coated with RGD peptides and imaged after 48 hours of exposure. Green = actin, Cyan = DNA, pink = bead. Scale bar = 10 µm.

**Figure S2.**
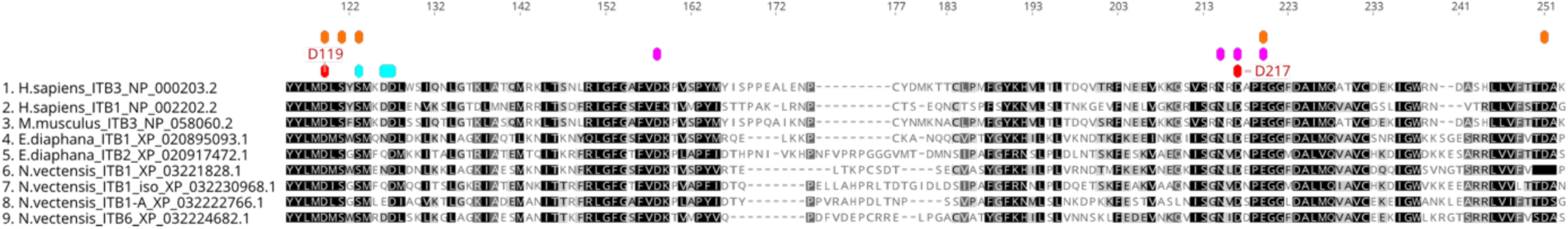
Active sites of RGD-binding integrin and mutations used for this study. Highly conserved residues are highlighted in black and as conservation decreases this goes from grey to white. Conserved active sites are noted above with the colored labels. Orange = MIDAS. Turquoise = ADMIDAS. Pink = LIMBS. Red = used mutations to disrupt RGD binding.

**Figure S3.**
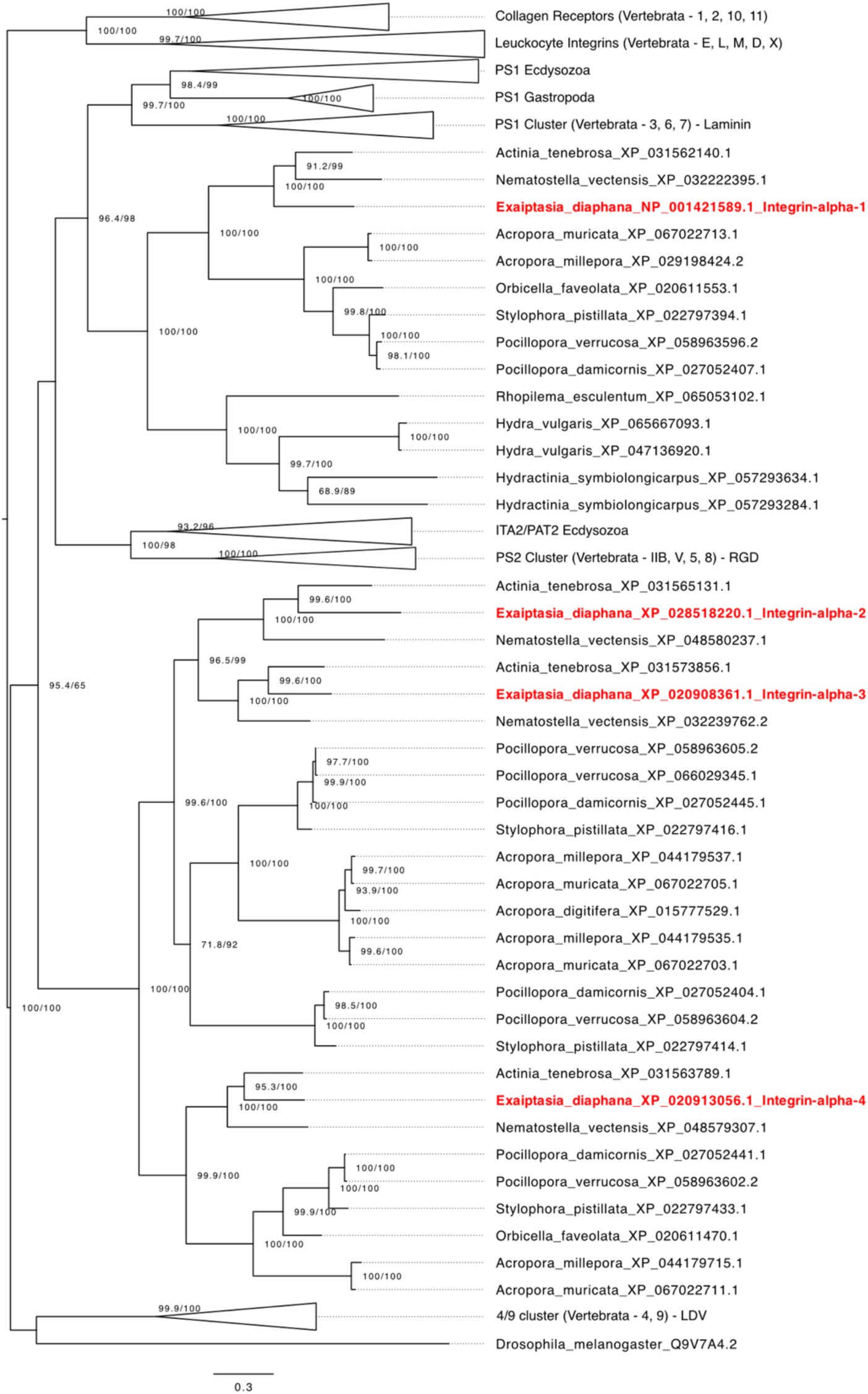
Phylogenetic tree of integrin ⍺ proteins prior to collapse of subtrees and rotation of nodes.

